# A primate specific loss of function polymorphism in TLR2 gene decreases inflammation and protects humans from organ dysfunction in Malaria

**DOI:** 10.1101/239129

**Authors:** Aditya K Panda, Ratnadeep Mukherjee, Bidyut K Das, Rina Tripathy, Ashok K Satapathy, Shobhona Sharma, Balachandran Ravindran

**Author notes:** Corresponding author: Balachandran Ravindran, Ph.D., Infectious Diseases Biology Laboratory, Institute of Life Sciences, Bhubaneswar- 751 023 India. Current address: Centre for Life Sciences, Central University of Jharkhand, Brambe, Ranchi, Jharkhand, India (Aditya K Panda). Current address: National Cancer Institute, NIH, Bethesda, MD, USA.

## Abstract

Polymorphisms of TLR genes could regulate and contribute functionally to innate immunity and inflammation. TLR2, a promiscous receptor recognizes Pathogen Associated Molecular Patterns from several microbes, bacterial, viral, protozoan and helminths. We demonstrate that monocytes of humans with deletion polymorphism in TLR2 gene (a 23 bp deletion in 5’ UTR region) respond more vigorously *in vitro* to several TLR2 ligands in comparison to those with insertion allele. Lower primates such as Rhesus monkeys and Baboon display ‘deletion’ genotype while insertion is found in higher primates viz., Orangutan, Chimpanzees and Gorilla. Enhanced inflammation is a hallmark of pathogenesis in human severe malaria leading to bad prognosis and odds ratio of patients prone to develop severe malaira such as multi organ failure with del/del genotype was found to be very high. Based on induction of inflammatory cytokines by normal human PBMCs *in vitro* and circulating cytokine levels in cohorts of patients with severe *P. falciparum* malaria, we propose that ‘insertion’ of a 23bp sequence in 5’UTR region of TLR2 gene could have led to moderated TLR2 induced inflammation thus offering survival advantage to higher primates by rendering them relatively refractory to multi-organ dysfunction in severe malaria.

**One Sentence Summary:** A 23bp deletion in TLR2 gene is associated with high inflammation and susceptibility to organ dysfunction in *Plasmodium falciparum* malaria.

## Introduction

Of all the mammalian TLRs characterized so far, TLR2 displays a high degree of promiscuity since PAMPs from several bacterial, viral, fungal, protozoan and helminthic pathogens, and DAMPs such as HMGB1, HSP60, HSP70 etc. stimulate through TLR2 resulting in inflammatory activation of macrophages, the cellular basis of both innate immunity and inflammation (*1, 2*). Polymorphisms in TLR2 gene thus determine clinical outcome of several human diseases (*3*). A 23 bp insertion/deletion polymorphism in 5’ UTR region of TLR2 gene reported in human populations and submitted by us to dbSNP (ss181129011 https://www.ncbi.nlm.nih.gov/projects/SNP/snp_ss.cgi?ss=181129011) was of specific interest to us for two reasons – a) the insertion was located in chromosome 4, approximately 60 bp downstream of an NF-kB binding site, a key region regulating inflammation (*4*) and b) although published literature describes this polymorphism as Insertion vs Deletion in several ethnic human populations – Ins allele being the dominant one, a search of database revealed that the 23bp insertion in the 5’ UTR region of TLR2 gene was unique and found only in primates and not in rodents and other mammals. More interestingly, in lower primates such as rhesus, nomascus monkeys, baboons etc insertion of one copy of the 23bp sequence was observed and in higher primates viz., Orangutans and Chimpanzees two copies were found to be inserted while in Gorilla three copies were found (summarized in Figure 1a). In the present study we set out to a) validate these insertion polymorphism in wild primates, and b) test in vitro response of human mononuclear cells with Indel polymorphism on stimulation with several TLR2 ligands, and c) to study the status of inflammatory response in patients with severe malaria to correlate the Indel polymorphism with disease outcome.

**Figure-1.**
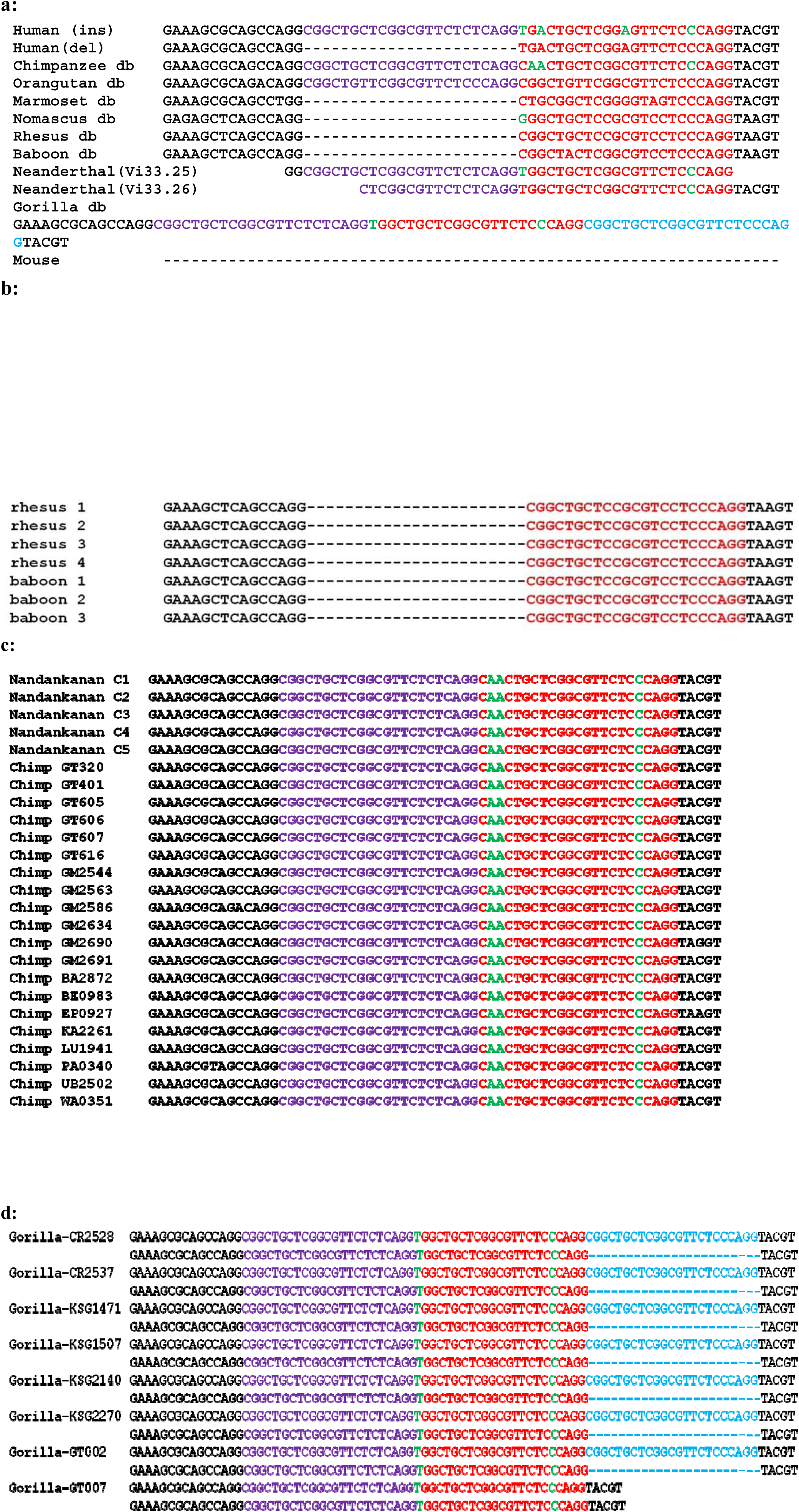
Multiple sequence alignment of 1st exon flanking TLR2 (23bp ins/del) polymorphism site in primates. **(a)**5’ UTR sequences in Human, Chimpanzee, marmoset, rhesus monkey, baboon were obtained from NCBI data base and Gorilla sequence from Ensembl genome browser. Neanderthal sequence was obtained from whole genome sequence database. The alignment was performed by web-based multiple sequence alignment programmer (CLUSTAL 2.0.12). Neanderthal (VI 33.25, VI 33.26) and gorilla sequences were aligned manually. Results revealed that Marmoset, Rhesus monkey and Baboon harbored del allele, and chimpanzee, orangutan and Neanderthals displayed ins allele. Gorilla TLR2 5’UTR displayed a double insertion sequence **(b)** To validate the above sequence in database, TLR2 5’UTR region was amplified using genomic DNA of 4 Rhesus monkeys, 3 Baboons, 25 Chimpanzees and 8 Gorilla samples. The amplicons were sequenced and aligned by web based multiple sequence alignment programmer. Lower primates Rhesus monkeys and Baboon displayed homozygous deletion allele, while higher primates viz. chimpanzee harbored insertion of 23bp nucleotides in both the strands. Out of eight gorilla sequenced for TLR2 Indel polymorphism, seven were heterozygous for double insertion type and one was homozygous for single insertion similar to twenty five chimpanzees investigated in this study.

## Results

We tested the 23 bp TLR2 polymorphism in 4 rhesus monkeys, 3 baboons, 25 Chimpanzees and 8 Gorilla using fecal DNA samples to validate gene sequences reported in NCBI data-base - polymorphism observed by us was consistent with sequences reported in the database – all the 25 Chimpanzees were homozygous for ‘del’ allele and 1 of the 8 Gorilla was heterozygous while the other 7 displayed homozygous double ‘Insertion’ (Figures 1b, 1c and 1d). A closer examination with rodent TLR2 gene sequence revealed that expression of 23bp ‘insertion’ and ‘deletion’ widely being reported by investigators in literature in humans and in the data base is a misnomer – the polymorphism is essentially absent in TLR2 5’ UTR region of rodent genes and lower primates seems to have acquired one copy since downstream of the 23bp ‘insertion’ a similar 23 bp sequence is found in rhesus monkeys, nomascus monkeys and baboon which are otherwise classified as ‘deletion’ in literature (fig 1a). Thus what has been designated as ‘deletion’ in lower primates in reality is ‘insertion’ when compared with 5’ UTR region gene of mice TLR2. In higher primates such as Chimpanzee and Organgutan the 23 bp sequence is found duplicated once and in Gorilla TLR2 gene it is duplicated twice. The ‘insertion’ event acquired by lower primates appears to have been positively selected during evolution. In this manuscript for the sake of clarity with current literature we continue to use the terminology being widely used currently viz., ‘deletion’ and ‘insertion’ of 23bp sequence of TLR2 gene.

Since wild life regulations do not allow undertaking phenotypic studies using cells of higher primates, we studied functional significance of this TLR2 polymorphism in humans with ‘del’ and ‘ins’ alleles. In a cohort of 432 normal healthy Indian adult population prevalence of ‘del’ allele was found to be about 14% - the frequency of ‘del’ polymorphism in other geographical areas varied from 13% in PNG to 36% in Northern Han Chinese. (Table 1). Normal healthy human volunteers with ‘del’ allele, expressed significantly more surface TLR2 on monocytes in comparison to those with ‘ins’ allele (Fig 2a) and also displayed elevated plasma TNF-α and CRP levels, (although at physiologically normal levels) markers of systemic inflammation (Fig 2b & 2c). A significant positive correlation was found between plasma levels of TNF-α and CRP (Fig 2d). These findings suggested a positive association between ‘del’ allele of TLR2 gene and biomarkers of systemic inflammation even in normal healthy adults. We tested the significance of this observation *in vitro* by directly testing for activation with TLR2 lignads. We stimulated normal human PBMCs with Pam3CSK4 and Vi polysaccharide of *Salmonella typhi* (*5*) and quantified cytokines in culture supernatants. Subjects with ‘del’ allele secreted higher levels of IL-6, IL-1β, TNF-α and IL10 in comparison to those homozygous for ‘ins’ polymorphism (Figure 3 a, b, c, d). There was no significant difference between the two groups when stimulated with CpG DNA, a TLR9 ligand. For further confirmation a larger panel of TLR2 ligands was used for stimulation of normal PBMCs in vitro. About 4 to 7 fold enhanced secretion of IL-1β was observed in subjects with ‘del’ allele when stimulated with not only with Pam3CSK4 and Vi polysaccharide but also with peptidoglycan (PGN) from 6 different microbial source or γ-irradiated *Mycobaterium tuberculosis* (Figure 3e), suggesting propensity of ‘del’ subjects to respond more vigorously to TLR2 ligands. The observed difference was specific to only TLR2 since TLR4, TLR5 and TLR9 expression were comparable in all the subjects (data not shown) and levels of IL-1b secretion was similar in both the groups when stimulated with poly IC, flagellin or CpG DNA (Figure 2e). We conclude that normal humans with ‘del’ allele in TLR2 gene display higher levels of plasma TNF-α and CRP and their PBMCs get activated more vigorously when stimulated with TLR2 ligands.

**Table-1.**
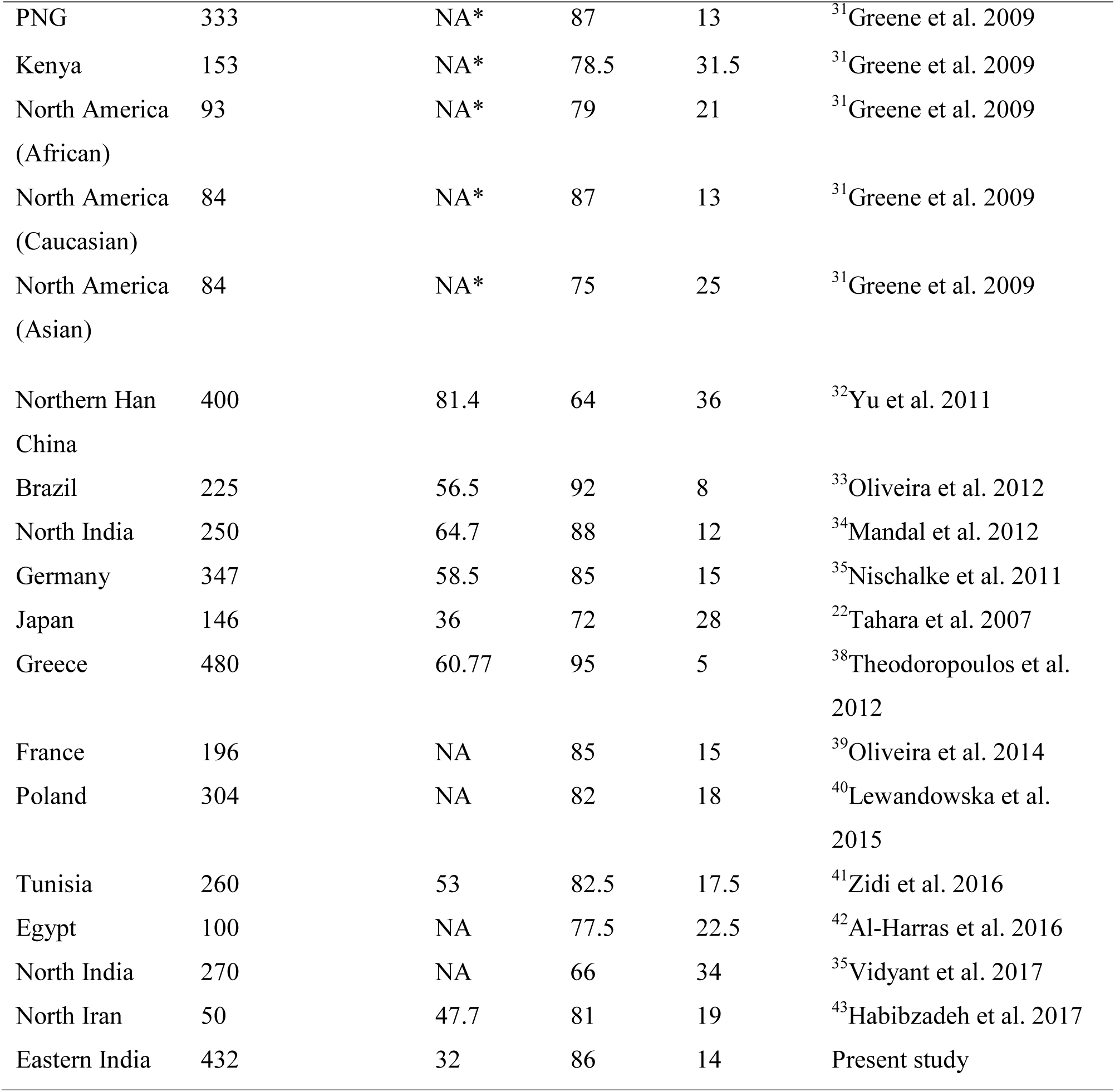
Distribution of TLR2 23bp indel polymorphism in various populations

| Studied area | Number of healthy subjects | Mean age (years) | ins allele (%) | del allele (%) | Reference |
| --- | --- | --- | --- | --- | --- |
| PNG | 333 | NA* | 87 | 13 | <sup>31</sup>Greene et al. 2009 |
| Kenya | 153 | NA* | 78.5 | 31.5 | <sup>31</sup>Greene et al. 2009 |
| North America<br>(African) | 93 | NA* | 79 | 21 | <sup>31</sup>Greene et al. 2009 |
| North America<br>(Caucasian) | 84 | NA* | 87 | 13 | <sup>31</sup>Greene et al. 2009 |
| North America<br>(Asian) | 84 | NA* | 75 | 25 | <sup>31</sup>Greene et al. 2009 |
| Northern Han<br>China | 400 | 81.4 | 64 | 36 | <sup>32</sup>Yu et al. 2011 |
| Brazil | 225 | 56.5 | 92 | 8 | <sup>33</sup>Oliveira et al. 2012 |
| North India | 250 | 64.7 | 88 | 12 | <sup>34</sup>Mandal et al. 2012 |
| Germany | 347 | 58.5 | 85 | 15 | <sup>35</sup>Nischalke et al. 2011 |
| Japan | 146 | 36 | 72 | 28 | <sup>22</sup>Tahara et al. 2007 |
| Greece | 480 | 60.77 | 95 | 5 | <sup>38</sup>Theodoropoulos et al.<br>2012 |
| France | 196 | NA | 85 | 15 | <sup>39</sup>Oliveira et al. 2014 |
| Poland | 304 | NA | 82 | 18 | <sup>40</sup>Lewandowska et al.<br>2015 |
| Tunisia | 260 | 53 | 82.5 | 17.5 | <sup>41</sup>Zidi et al. 2016 |
| Egypt | 100 | NA | 77.5 | 22.5 | <sup>42</sup>Al-Harras et al. 2016 |
| North India | 270 | NA | 66 | 34 | <sup>35</sup>Vidyant et al. 2017 |
| North Iran | 50 | 47.7 | 81 | 19 | <sup>43</sup>Habibzadeh et al. 2017 |
| Eastern India | 432 | 32 | 86 | 14 | Present study |

**Figure-2.**
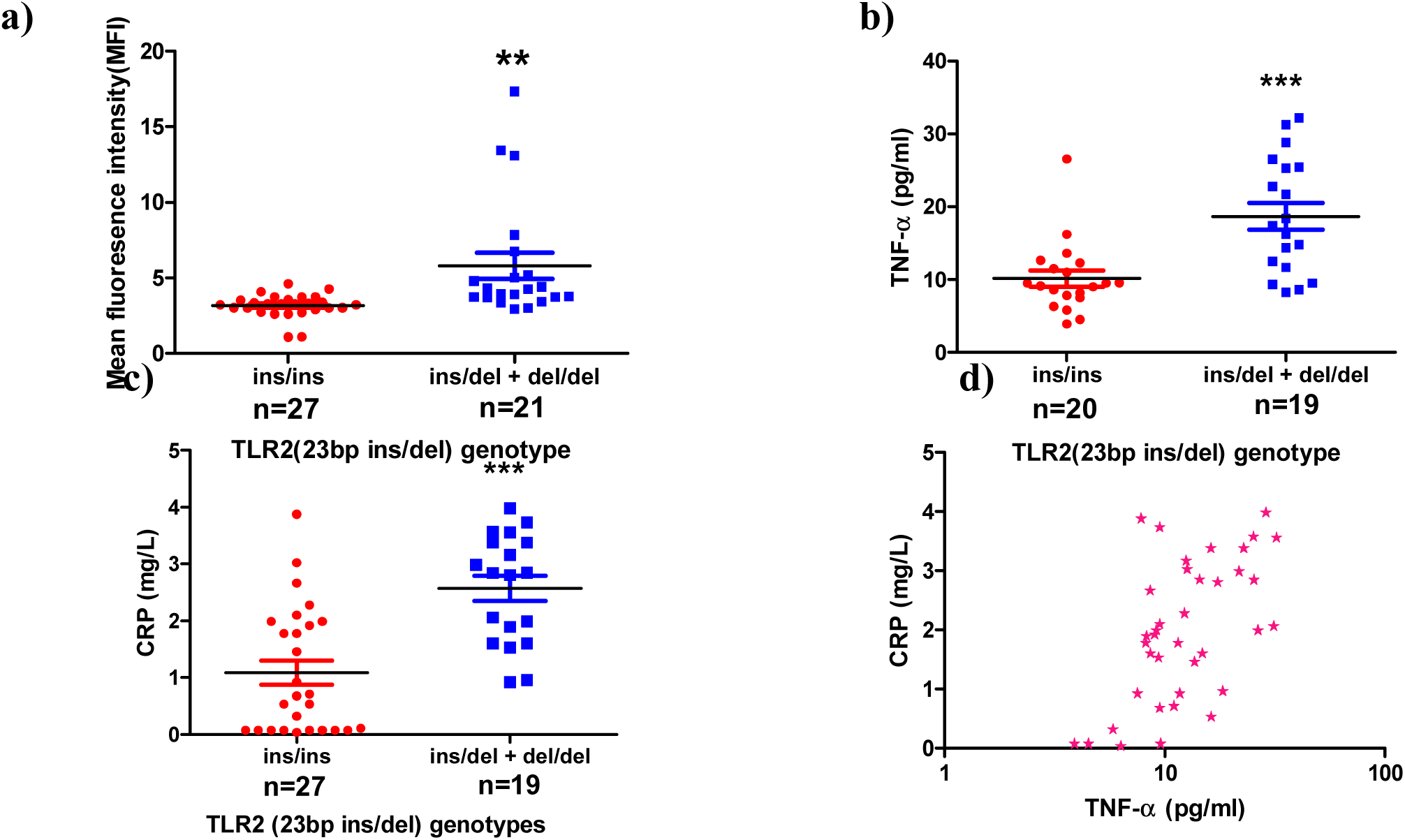
TLR2 (23bp ins/del) genotype is associated with higher TLR2 expression and plasma markers of inflammation: PBMCs of healthy laboratory volunteers were stained for TLR2 and CD14+ monocytes were analyzed by flow cytometry; TNF-α and CRP were quantified in their plasma. **a:** Mean fluorescence intensity (MFI) of surface TLR2 expression after normalization with isotype control. TLR2 expression was significantly more (P<0.0001) in subjects with ‘del’ allele in comparison to those with ‘ins’ allele. Plasma levels of TNF-α **(b)** and CRP **(c)** were quantified by ELISA and a turbidolatex method respectively. Significantly higher levels of both plasma TNF-α (P<0.001) and CRP (unpaired t test P<0.001) was observed in ‘del’ carriers; **(d)** A significant positive correlation was demonstrable between plasma CRP and TNF-α levels in 46 healthy controls (P<0.0001, Spearman r=0.53). Significance in figure 1a remained even when the three out layers were not factored-in for statistical analysis.

**Figure-3.**
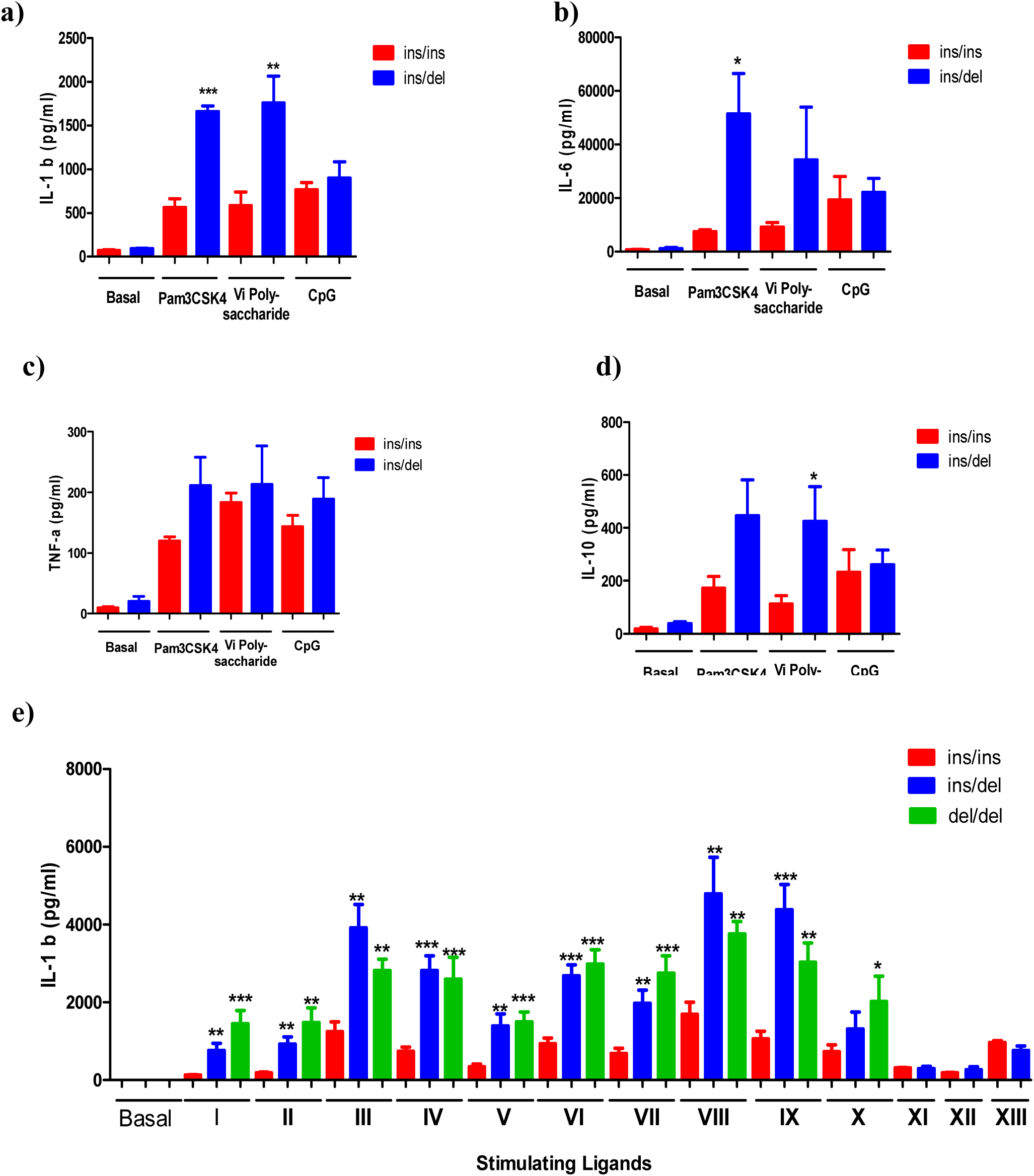
TLR2 ligands induce enhanced levels of cytokines in PMBCs of ‘del’ allele carriers: Healthy controls were screened for TLR2 (23bp ins/del), TLR4 (Asp299Gly and Thr399Ile) and TLR1 (I602S) polymorphisms as described in methods section. Individuals, with wildtype TLR4 and TLR1 polymorphisms i.e. Asp299Asp, Thr399Thr and I602I but displaying ‘del’ or ‘ins’ of TLR2 were tested. Normal human PBMCs with ins/ins (n=5) or ins/del (n=5) were stimulated in separate matrixes with Pam3CSK4 (1ng/ml), Vi polysaccharide (20ug/ml), CpG (10ug/ml) or LPS (10ug/ml) for 48hrs. Polymixin B (10ug/ml) was used in culture medium except in positive controls where LPS was employed as a ligand. In all assays LPS with polymixin-B was taken as negative control to ensure non-induction of IL1β. Culture supernatants were quantified for **(a)** IL-1b, **(b)** IL-6, **(c)** TNF-α and **(d)** IL-10 by Bio-plex bead array (Biorad). Upon stimulation with Pam3CSK44, cells of ins/del subjects produced significantly higher IL-1b (P<0.001) and IL-6 (P<0.05) when compared to ins/ins subjects; Vi polysaccharide of S.typhi induced higher IL-1b (P<0.01) and IL-10 (P<0.05) in ins/del carriers. CpG DNA a TLR-9 ligand induced comparable levels of four cytokines in both the groups of subjects; **(e)**, human PBMCs from 8 ins/ins, 8 ins/del 5 and del/del subjects were stimulated with (I) Pam3CSK44 (1ng/ml), (II) Vi polysaccharide (20ug/ml), (III) Peptidoglycan (PGN) from Streptomyces (100ug/ml), (IV) PGN from Methano species (100ug/ml), (V) PGN from Saccharomyces cereviseae (100ug/ml), (VI) PGN from Micrococcus luteus (100ug/ml), (VII) PGN from Bacillus subtilis (100ug/ml), (VIII) PGN from Staphylococcus aureus (100ug/ml) and (IX) γ- irradiated Mycobacterium tuberculosis (1X10^6^ /ml). (X): LPS (10ug/ml) was taken as positive control. As controls PBMCs from 6 ins/ins and 6 ins/del carriers were stimulated with (XI) CpG (10ug/ml), (XII) Poly IC (10ug/ml) and (XIII) Flagellin (100ug/ml). IL-1b levels were measured in all culture supernatants by ELISA and analyzed by student’s t test. P value less than 0.05 was considered as significant (* = P < 0.05, ** = P < 0.01 and *** = P< 0.001). Subjects with ‘del’ allele consistently secreted significantly higher levels of IL-1b when stimulated with any of the TLR2 ligands while supernatants of cultures stimulated with CpG, Poly IC and Flagellin displayed comparable levels of IL-1b. Polymyxin-B at a concentration of 10ug/ml completely inhibited LPS mediated activation (which was taken in each plate) - IL-1b levels below 10pg/ml in LPS stimulated cavities (with Polymyxin-B) were treated as invalid experiments.

Since the insertion of 23bp sequence in TLR gene is observed only primates and not in lower animals such as rodents we addressed the issue of significance of this insertion during primate evolution. ‘Malaria hypothesis’ widely reported in literature attributes severe *P. falciparum* for selection of a variety of haemoglobinopathies reported in human population (*6*). Secondly, malarial glycolsyl-phosphatidylinositol (GPI), a TLR2 ligand has been proposed as a ‘malaria toxin’ that induces inflammatory activation of monocytes/macrophages and is reported to mediate systemic inflammation and play a role in mediating severe clinical manifestations observed in malaria (*7, 8*). Based on this logic malarial GPI has also been tested as a candidate anti-disease vaccine for malaria (*9*). We hypothesized that malaria could have offered survival advantage in patients with insertion allele of 23bp sequence in their TLR2 gene that leads to decreased inflammatory response thus contributing to lower inflammatory response, lesser clinical severity and mortality caused by malaria. To test our hypothesis we enrolled 398 malaria patients with different grades of clinical severity viz., un-complicated malaria (UM), cerebral malaria (CM), multi organ dysfunction (MOD) and non-cerebral severe malaria (NCSM) (*10*) (Table 2) and genotyped TLR2 ins/del polymorphism in 356 patients (Table 3). Patients with ‘del’ polymorphism displayed predilection for development of severe malaria (Table 3) and increased mortality (Table 4). Plasma TNF-α and CRP levels correlated positively with clinical severity (Fig 4a and 4b) as well as mortality (Fig 4c and 4d) in the cohort of malaria patients analyzed in the study, confirming earlier reports of direct correlation between increased inflammation and bad prognosis in malaria (*11, 12*). More crucially, increased plasma TNF-α and CRP levels in our patients correlated very significantly with TLR2 genotype - patients with ‘del’ allele were found to be high producers of TNF-α and CRP (Fig 5a and 5b respectively) in comparison to those homozygous for ‘ins’ polymorphism, suggesting a significant and association between this TLR2 polymorphism and enhanced systemic inflammation and clinical severity in human *P. falciparum* malaria. This association was found to be consistent when each of the clinical groups were analyzed for plasma TNF-α (Fig 5c respectively). Polymorphism of TNF-α genes and its association with enhanced TNF-α levels and clinical severity have been reported (*13*) – in comparison our observations on TLR2 ‘del’ polymorphism in adult population and its association with plasma TNF-α levels as well as clinical severity and mortality appears to be most compelling. Another polymorphism viz., GT repeat microsatellite in intron2 of TLR2 gene has been earlier shown to determine TLR2 mediated inflammation in humans - larger the number of GT repeats lesser is the inflammatory response to TLR2 ligands (*14*, *15*). Such GT repeats are absent in lower primates and higher primates display GT repeats (*16*). We typed our cohort of malaria patients to investigate possible relationship between GT repeat polymorphism and plasma TNF-α and CRP levels. There was no significant association between GT repeat polymorphism and plasma levels of TNF-α and CRP (Fig 6a and 6b respectively). When TLR2 23bp polymorphism and GT repeats were combined and analyzed together, patients with ‘del’ allele were found to have significantly more TNF-α and CRP in patients regardless of short or long GT repeats in TLR2 gene (Fig 6c and 6d respectively)

**Table-2.** Characteristic of *P. falciparum* infected Indian patients at hospital admission.

**Table-1 Distribution of TLR2 23bp indel polymorphism in various populations** **Table-2 Characteristic of *P. falciparum* infected Indian patients at hospital admission.**
| Characteristics | CM (n=97) | MOD (n=123) | NCSM (n=75) | UCM (n=103) |
| --- | --- | --- | --- | --- |
| Sex (M/F) | 73/24 | 97/26 | 60/15 | 79/24 |
| Mean age in years (range) | 30.77 (15-86) | 32.09 (15-83) | 33.02 (15-70) | 31.19 (15-75) |
| Hemoglobin g/dl, mean ± SD | 9.879 ± 2.028 | 9.848 ± 2.206 | 9.704 ± 2.488 | 10.54 ± 1.867 |
| GCS (mean (range)) | 8.598 (3-11) | 7.98 (3-15)* | -- | -- |
| Neutrophils%,mean(range) | 68.52 (42-92) | 70.2 (37-92) | 67.64 (31-88) | 65.88 (33-92) |
| Lymphocytes%,mean(range) | 27.15 (10-54) | 26.33 (5-59) | 27.10 (0-65) | 29.32 (7-55) |
| Monocytes %, mean (range) | 1.57 (0-4) | 1.58 (0-4) | 1.38 (0-5) | 1.21 (0-6) |
| Eosinophils %, mean (range) | 3.11 (0-12) | 2.82 (1-7) | 3.19 (2-16) | 3.17 (0-10) |
| Total leukocyte count, mean ± SD | 8431 ± 2097 | 9299 ± 2875 | 8512 ± 2433 | 7850 ± 2162 |
| TPC mean ± SD | 188333 ± 52915 | 178625 ± 67712 | 174720 ± 74581 | 197267 ± 64898 |
| Fasting blood sugar mg/dl, mean ± SD | 148 ± 84.21 | 141.7 ± 98.34 | 138.5 ± 82.40 | 120.7 ± 56.0 |
| SGOT units/ml, m ± SD | 30.13 ± 7.99 | 169.9 ± 120.7** | 130.8 ± 78.19** | 33.96 ± 8.48 |
| SGPT units/ml, m ± SD | 27.47 ± 8.94 | 130.5 ± 75.48** | 109.5 ± 54.28** | 28.08 ± 9.97 |
| ALP, mean ± SD | 170.8 ± 93.16 | 221.7 ± 156.6 | 190 ± 141.7 | 158.3 ± 82.2 |
| Serum urea mg/dl, m ± SD | 53.31 ± 28.82 | 142.5 ± 71.85*** | 169.3 ± 73.00*** | 37.73 ± 19.04 |
| Serum creatinine mg/dl, mean ± SD | 1.36 ± 0.63 | 5.03 ± 3.38*** | 6.13 ± 3.05*** | 1.15 ± 0.34 |
| Na<sup>+</sup> mEq/L, mean ± SD | 136.2 ± 8.07 | 134.4 ± 9.51 | 133.3 ± 6.47 | 135.9 ± 7.96 |
| K<sup>+</sup> mEq/L , mean ± SD | 4.3 ± 0.91 | 4.51 ± 1.12 | 4.21 ± 0.8 | 4.17 ± 0.73 |
| Mg<sup>++</sup> mEq/L, mean ± SD | 3.48 ± 2.06 | 3.27 ± 1.34 | 3.34 ± 1.58 | 3.52 ± 3.30 |
| Ca<sup>++</sup> mEq/L, mean ± SD | 0.87 ± 0.20 | 0.99 ± 0.15 | 0.89 ± 0.19 | 0.92 ± 0.21 |
| Case fatality rate, no (%) | 13 (14) | 36 (32) | 0 | 0 |

Note: \*CM+MOD, \*\*Hepatopathy, \*\*\*Acute renal failure

**Table 3.**
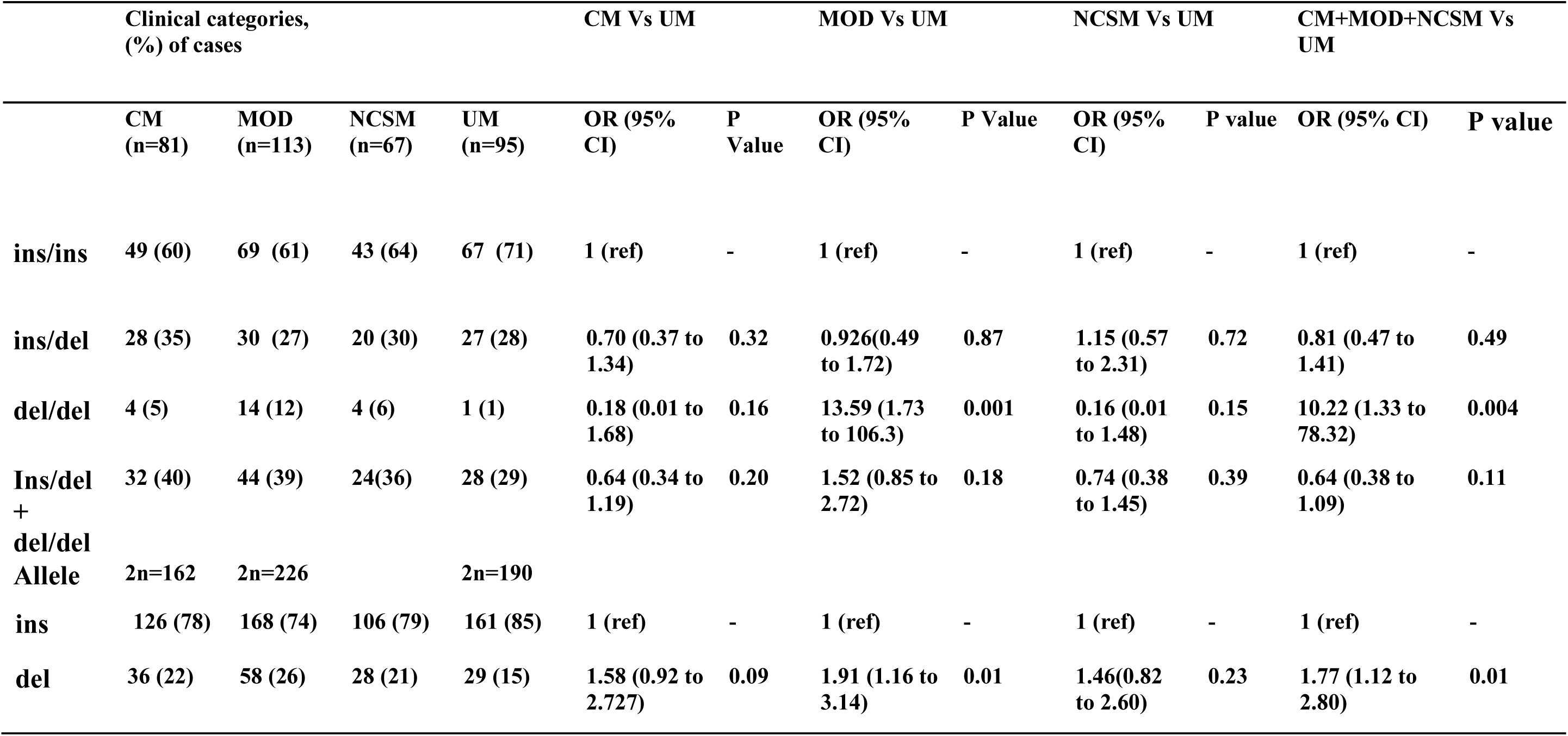
Prevalence of Genotype and allele of TLR2 (23bp ins/del) polymorphism in P.falciparum malaria patients.

**Table-4.** Association of TLR2 (23bp ins/del) polymorphism with treatment out come in severe *P. falciparum* malaria.

| **Genotype/Allele** | **Death** | **Survivor** | **P value** | **OR (95% CI)** |
| --- | --- | --- | --- | --- |
| **Genotype** | **(n=36)** | **(n=158)** |  |  |
| ins/ins | 16 (44) | 102 (65) | - | 1 (ref) |
| ins/del | 13 (36) | 45 (28) | 0.19 | 1.84 (0.81 to 4.14) |
| del/del | 7 (20) | 11 (7) | 0.01 | 4.05 (1.37 to 12.00) |
| ins/del + del/del | 20 (56) | 56 (35) | 0.03 | 2.27 (1.09 to 4.74) |
| **Allele** | **(2n=72)** | **(2n=316)** |  |  |
| ins | 45 (63) | 249 (79) |  |  |
| del | 27 (37) | 67 (21) | 0.005 | 2.23 (1.28 to 3.85) |

**NOTE: Data are no. (%) of participants unless otherwise specified;**
**OR: odds ratio; CI: confidence interval**

**Figure-4.**
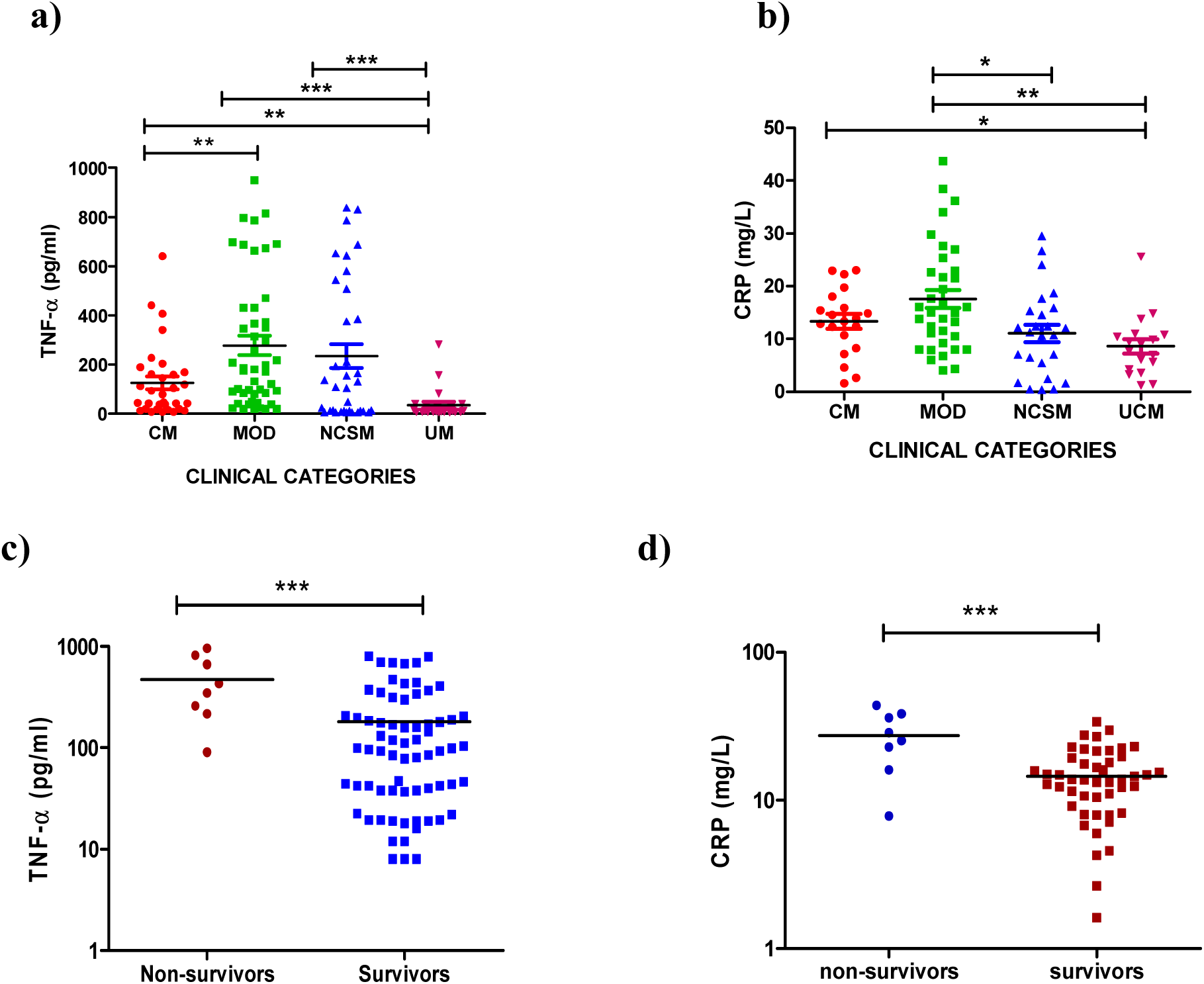
Plasma TNF-α and CRP levels correlate with clinical severity and mortality in malaria. **(a,b):** Plasma TNF-αand CRP levels were determined by ELISA (CM, n=32; MOD, n=47; NCSM, n=35 and UM, n=24) and turbidometric assays (CM, n=20; MOD, n=35; NCSM, n=25 and UM, n=18), respectively in samples collected from patients before treatment with anti-malarials. CM and MOD cases were enrolled for correlation between TNF-α and CRP levels with mortality. Plasma levels of TNF-α (**c**) and CRP (**d**) was significantly more in patients who died when compared with survivors. (*P<0.05, **P<0.01, ***P<0.001)

**Figure-5.**
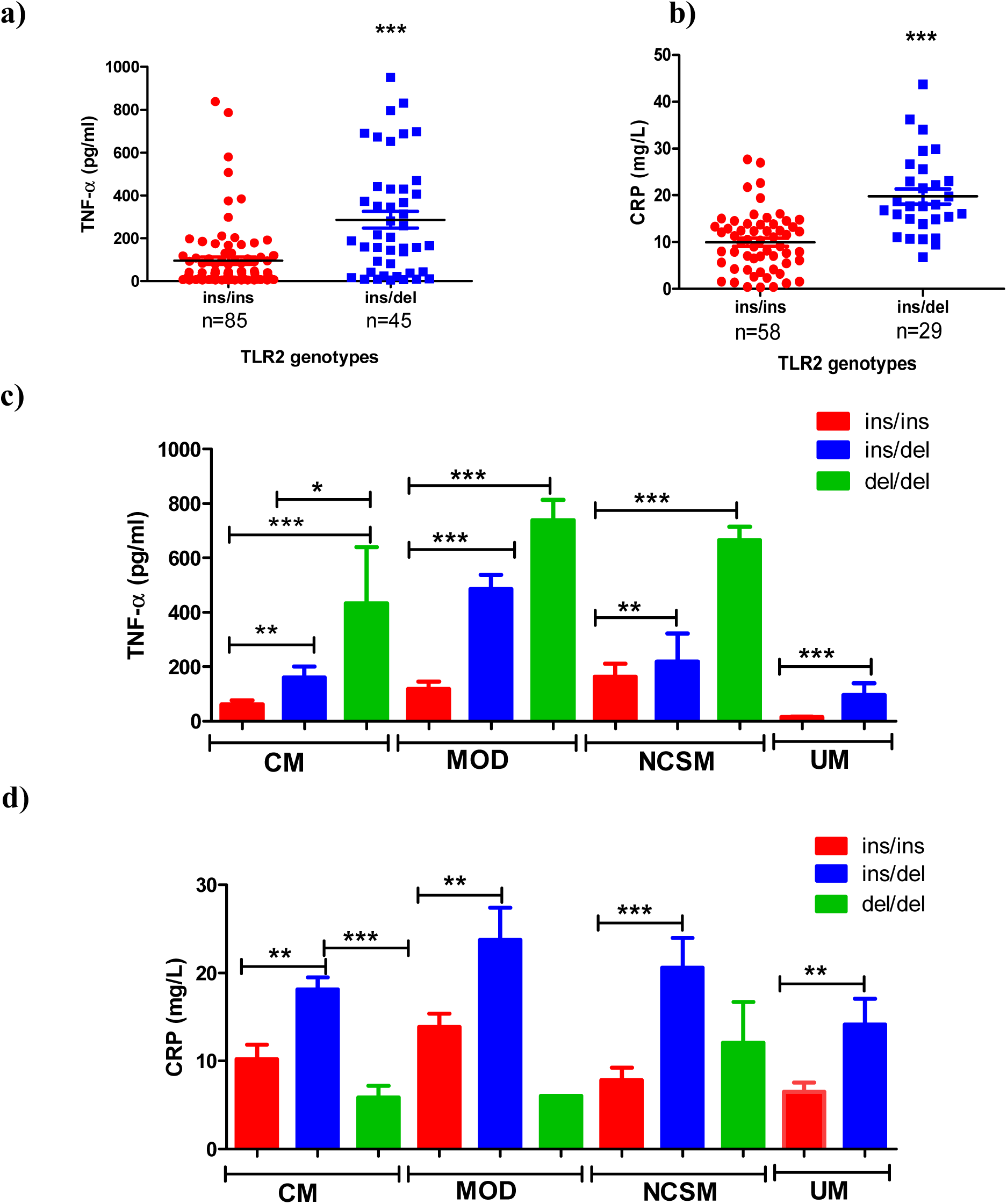
TLR2 ‘del’ allele is associated with higher plasma levels of TNF- α and CRP in *P. falciparum* infected patients. Plasma concentrations of **(a)** TNF-α and **(b)** CRP were measured by ELISA and Turbidolatex, respectively in plasma of *P. falciparum* infected patients (combined clinical categories) and correlated with ‘del’ or ‘ins’ genotype of TLR2 gene. Scatter plots show data for TNF-α in 130 patients and for CRP in 87 patients. Mean plasma levels of TNF-α and CRP were significantly elevated in malaria patients with ins/del genotype in comparison to patients with ins/ins (unpaired t test, ***P<0.001).**(c)** Plasma TNF-α levels in clinical categories of *P. falciparum* malaria viz. CM, cerebral malaria (ins/ins=17; ins/del=13; del/del=2); MOD, multi organ dysfunction (ins/ins=28; ins/del=17; del/del=2); NCSM, non-cerebral severe malaria (ins/ins=22; ins/del=9; del/del=4); UM, uncomplicated malaria (ins/ins=18; ins/del=6) were analysed by student’s t test in various genotypes.(*P<0.05, **P<0.01, ***P<0.001) (d) Plasma levels of CRP was compared different TLR2 indel genotypes in various clinical categories of *P. falciparum* malaria (**P<0.01,***P<0.001).

**Figure- 6.**
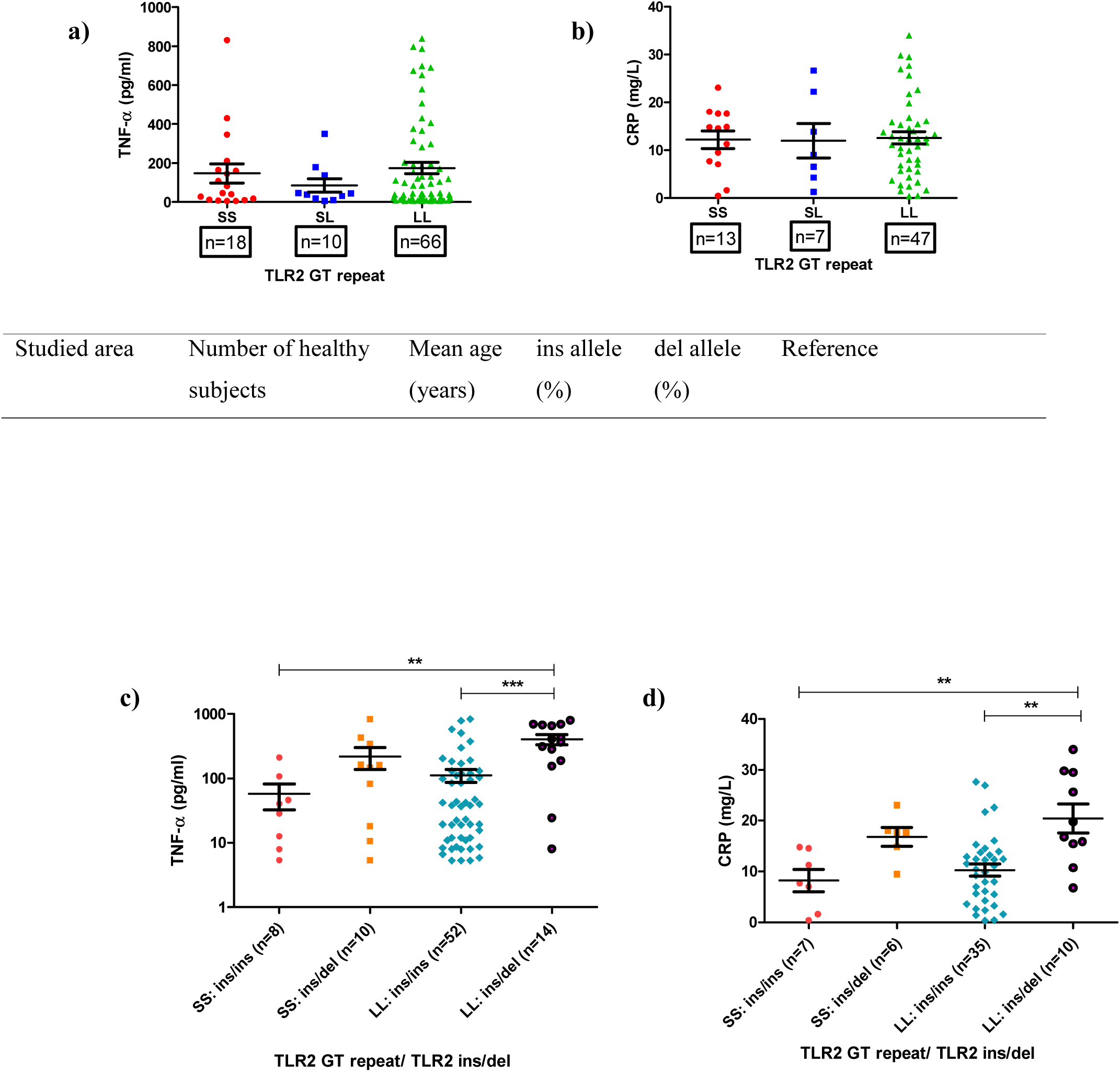
Plasma TNF- α and CRP levels are not correlated TLR2 (GT repeat) in P. *falciparum* malaria patients. TLR2 GT repeat was typed by PCR-fragment analysis. Alleles were classified in to two broad group S (<11 repeat) and L (16-21 repeats). Plasma concentrations of TNF-α and CRP were measured by ELISA and Turbidolatex, respectively in plasma of *P. falciparum* infected patients (combined clinical categories) and correlated with GT repeat alleles (S and L) of TLR2 gene. Mean plasma levels of TNF-α and CRP were comparable (**a** and **b**). On combined analysis of TLR2 (23bp ins/del) and TLR2 GT repeat polymorphism (**c** and **d**), patients with LL: ins/del genotypes combination displayed higher plasma TNF-α and CRP when compared to SS: ins/ins and LL: ins/ins combinations. (*P<0.05, **P<0.01, ***P<0.001)

## Discussion

The current study has offered several novel insights in the context of a TLR-2 polymorphism, evolution of TLR-2 in primates and susceptibility to TLR2 mediated inflammation in human malaria. First, ‘deletion’ of a 23bp nucleotide sequence in TLR-2 observed only in lower primates (Rhesus monkey, Baboon, Marmoset and Nomascus) and not in higher primates (Chimpanzees, Gorilla, Orangutan and Neanderthal) as reported in literature appears to be a misnomer - in comparison with TLR2 gene of rodents, the 23bp sequence was indeed found to be already inserted in 5’ UTR region in lower primates that are otherwise designated as ‘deletion’ in literature. The ‘insertion’ seems to have been acquired by lower primates and positively selected during evolution that appears to have been duplicated further in higher primates viz., Chimpanzees, Orangutan and Gorilla. Distribution pattern of TLR2 23bp ins/del polymorphism as reported in database, were confirmed by us by typing polymorphism in 4 rhesus monkeys, 3 baboons, 8 Gorillas and 25 Chimpanzees collected from the wild. In the context of very high prevalence of asymptomatic malaria (analogous to asymptomatic infections in malaria endemic human population) in wild Chimpanzees and Gorilla (*17*) and two and three 23bp ‘insertion’ sequences respectively in Chimpanzee and Gorilla we propose that malaria may have offered evolutionary pressure to select ‘Ins’ allele of TLR2 gene in primates.

We sought evidence for this proposal by studying a large cohort of malaria patients since ‘Malaria hypothesis’ widely reported in literature attributes severe *P. falciparum* infections in human communities for selection of a variety of haemoglobinopathies in human population (*6*). Secondly, malarial glycosylphosphatidylinositol (GPI), a TLR2 ligand has been proposed as a ‘malaria toxin’ that induces inflammatory activation of monocytes/macrophages and responsible for severe clinical manifestations observed in malaria (*7, 8*). Based on this logic malarial GPI has also been tested as a candidate anti-disease vaccine for malaria (*9*). We hypothesized that severe malaria could have offered evolutionary pressure needed for selecting ‘ins’ allele in TLR2 gene in higher primates leading to decreased inflammation and contributing to lower clinical severity and mortality caused by *P. falciparum* malaria. Reports in literature offer credence to this view. Chimpanzees and Gorilla have restricted natural habitat with small population size and about 40-50% of wild chimpanzees and gorillas have been found infected (analogous to asymptomatic infection status in humans) with *P. falciparum* ‘like’ malarial parasites (*17, 18*). On the contrary, severe malaria such as cerebral malarial anemia, multi-organ failure etc are common cause of high mortality in human *P. falciparum* malaria (*19*) mediated by induction of inflammatory cytokine storm particularly high TNF-α levels leading to enhanced clinical severity and mortality (*6, 12*). To test our hypothesis we enrolled 398 malaria patients with grades of clinical severity viz., uncomplicated malaria (UM), cerebral malaria (CM), multi organ dysfunction (MOD) and non-cerebral severe malaria (NCSM) (*10*) and genotyped TLR2 ins/del polymorphism in 398 patients. Patients with ‘del’ allele displayed increased predilection for development of severe malaria and increased mortality. Inflammatory cytokine storm and high levels of plasma TNF-α have been attributed to development of pathology leading to bad prognosis in severe malaria (*6, 12*). In the cohort of malaria patients studied by us, we observed a significant positive association between high TNF-α levels and clinical severity: patients with MOD displayed the highest levels, followed by NCSM and CM. Plasma TNF-α levels were significantly more in patients who succumbed to severe malaria when compared to survivors, corroborating with earlier reports (*6, 12, 20*). While the role of very high levels of TNF-α in shock and organ failure leading to mortality has been known (*21*), the role of CRP in *P. falciparum* malaria is poorly documented in literature. CRP has been shown to bind to infected erythrocytes and clear damaged RBCs from circulation which possibly leads to anemia (*22*). Further, high levels of CRP has been related with parasite density and malarial severity (*23*). However the more significant and novel findings of the current study is association between elevated plasma levels of TNF-α and CRP with the TLR2 Indel genotype - patients with ‘del’ allele were found to be high producers of TNF-α and CRP in comparison to those with ‘ins’ allele, suggesting that patients with ‘deletion’ of 23bp sequence of TLR2 gene are prone to enhanced systemic inflammation leading to higher clinical severity when they get infected with *P. falciparum* malaria. Polymorphism of TNF-α genes and its association with enhanced TNF-α levels and clinical severity has been reported (*13*) – our observations on TLR2 ‘del’ polymorphism in adult population and its association with TNF-α levels as well as clinical severity and mortality appears most compelling. Since we found ‘ins’ allele consistently in higher primates in the wild (a large population of which harbor malarial parasites in circulation) and association of ‘del’ allele with increased clinical severity and mortality in humans with *P. falciparum*, we propose that malaria could have offered evolutionary pressure for selection of the 23 bp sequence in TLR-2 genes in higher primates thus offering them survival advantage.

While analyzing several wild primates we found consistent presence of ‘del’ allele in all lower primates and absence of ‘del’ allele in all Chimpanzees and Gorilla. It is not clear to us currently how about 20% of humans display ‘del’ allele in several ethnic human populations. The results do not however rule out complete absence of ‘del’ allele in higher primates since we have tested only a limited number of Chimpanzees and Gorilla that had survived selection under wild forest conditions. The other possibility is that presence of ‘del’ allele in about 20% of humans suggests that selection due to severe forms of malaria may have diminished in early humans due to absence of severe malaria leading to loss of ‘ins’ allele since it has been argued that emergence of severe *P. falciparum* malaria is a more recent evolutionary event (*24, 25*). It has been proposed that early homo sapiens lost their ability to synthesize N-Glucoryl neuraminic acid and consequently became resistant to *P. reichnowi* (malarial parasites infecting Chimpanzees) until emergence of *P. falciparum*, that uses N-Acetyl neuraminic acid as its receptor (*25*). Based on genetic analysis of several strains of *P. falciparum*, mosquito biting behavior and emergence of haemoglobin disorders selected by *P. falciparum* in Africa it has also been suggested that virulent *P. falciparum* may have emerged in human population only about 5000-10000 years ago as a consequence of introduction of agrarian practice (*24*).

The possibility that PBMCs with ‘del’ allele are prone to enhanced activation by TLR2 ligands was also tested. Stimulation of PBMCs of normal healthy volunteers with TLR-2 ligands such as PAM3CSK4, peptidoglycans from several bacterial species, Vi polysaccharide of *Salmonella typhi* etc revealed that subjects with ‘del’ allele respond more vigorously releasing significantly high levels of IL-1β, IL-6 and TNF-α when compared to those with ‘ins’ allele. These findings are in contrast to two earlier reports using experimentally transfected cell lines and scoring luciferase activity (*26, 27*) – we attribute this to possible experimental artifacts while using cell line models for studying functional activation of immune cells. Similar discrepancy between transfection-based functional analysis and in vivo response has been reported for TIRAP S180L polymorphism (*28*). We conclude that transfection-based experiments need to be interpreted with caution while correlating genotypes with phenotypes.

Another polymorphism in intron 2 region of TLR2 gene GT repeat microsatellite has been shown to determine TLR2 mediated inflammation in humans - larger the number of GT repeats lesser is the inflammatory response to TLR2 ligands (*15*). In the present study, no significant association of TLR2 GT repeats with plasma TNF-α and CRP was observed. Combined analysis of ‘del’ allele and GT repeat of TLR2 revealed that ‘del’ allele plays a more important role in induction of increased TNF-α and CRP levels in malaria than GT repeat polymorphism. More significantly, combined distribution of 23bp ins/del and GT repeat polymorphism reveled that larger GT repeats (reported to be less inflammatory) are linked to ‘ins’ allele. Our observation of absence of such GT repeats in lower primates and presence only in higher primates, while corroborating with a previous report (*16*) also supports the notion that decreased TLR2 mediated inflammation is linked to primate evolution. Regardless of malaria contributing to positive selection and emergence of ‘ins’ allele of TLR2 gene in higher primates, the observation that humans with ‘del’ allele are prone to increased TLR2 mediated inflammation has opened up avenues for future investigations in human innate immunity/inflammation. Both innate as well as adaptive immune responses are directly influenced by TLR-2 mediated activation of dendritic cells and T lymphocytes (*29–31*) and thus ‘ins’ vs ‘del’ in TLR2 gene could influence host response and susceptibility of humans to several other diseases such as HIV-AIDS (*32*), *H. pylori* mediated gastric cancer (*33*) and Alzheimer’s disease (*34*).

## Materials and Methods

### Ethics statement

Human blood samples from healthy controls (laboratory volunteers) and *P. falciparum* infected patients were collected using standardized Institutional protocol approved by Human ethical committee of Institute of Life Sciences, Bhubaneswar. Written informed consent was obtained from each of study participants or accompanying persons in the case of children and patients in comatose state.

### Healthy controls

123 healthy laboratory volunteers from Institute of Life Sciences and Regional Medical Research Centre, Bhubaneswar, India were enrolled. About 50ul of blood was collected by finger prick and genomic DNA was purified using a kit according to manufacturers’ instructions (Sigma). All individuals were genotyped for TLR2 (23bp ins/del), TLR2 codon (Arg677Trp, Arg753Gln), TLR4 (Asp299Gly, Thr399Ile) and TLR1 (I602S) polymorphisms as described below. To avoid confounders of activation by TLR2 ligands only those healthy subjects with wild type TLR4 and TLR1 polymorphism were used to investigate phenotype differences between 23bp Indel polymorphism. In addition, 309 healthy controls were enrolled and genotyped for TLR2 indel polymorphism.

### Detection of TLR2 (23bp ins/del, Arg753Gln and Arg677Trp) polymorphisms

All subjects were genotyped for TLR2 (23bp ins/del) polymorphism by polymerase chain reaction as described earlier (*35*). In brief, following primers were used for (23bp ins/del) genotyping: sense 5′-CACGGAGGCAGCGAGAAA-3’ and anti-sense 5′-CTGGGCCGTGCAAAGAAG-3’. PCR was carried out in a total reaction volume of 20 μL containing about 200 ng genomic DNA, 10 pmol each primer, 200 ng each dNTP and 1 U Taq DNA polymerase (sigma). The thermo-cycling condition was as follows: initial denaturation for 5 min at 95°C, followed by 35 cycles of 95°C for 30 s, 60°C for 30 s, and 72°C for 30 s. The final extension step was prolonged to 7 min at 72°C. PCR products were separated on 3% agarose gel electrophoresis and visualized in gel documentation system after ethidium bromide staining. A single band at 286 bp was classified as wild type (Ins/Ins), a single 263-bp band as homozygous mutation (Del/Del), and those displaying two bands of 286bp and 263bp were as heterozygous mutation (Ins/Del). Randomly chosen PCR products were purified using Qiagen PCR Purification System (Qiagen, UK) and sequenced using an ABI Prism 310 Genetic Analyzer (Applied Biosystems, Foster City, CA) and Big Dye Terminator DNA sequencing kit (Applied Biosystems). Distributions of two TLR2 polymorphisms (Arg753Gln, Arg677Trp) samples were genotyped by PCR followed by restriction fragment length polymorphism (RFLP) according to an earlier report (*36*). Primers: Forward 5′-GCCTACTGGGTGGAGAACCT-3′ and Reverse; 5′-GGCCACTCCAGGTAGGTCTT-3′were used to amplify a region of 340 bp including both polymorphisms (Arg753Gln, Arg677Trp) by following conditions: 940C for 5 min, then 30 cycles of 940C for 30 sec, 580C for 30 sec, and 72°C for 30 sec. The amplicons were digested overnight with *AciI* and analyzed by agarose gel electrophoresis. Amplified products were randomly sequenced for confirmation of individual’s genotype.

### Determination of TLR2, TLR4, TLR5 and TLR9 expression on circulating human monocytes

The TLR2 expression on cell surface was determined by flow cytometry using two-color immunofluorescent technique in 48 healthy laboratory volunteers: 27 ins/ins individuals, 16 subjects of ins/del and 5 subjects of del/del type were included for analysis. Briefly, 50ul of whole blood were directly stained for 30 min at 4 °C with a combination of fluorochrome-conjugated mAbs: phycoerythrin (PE)-labeled anti-TLR2 (Imgenix), fluorescein-isothiocyanate (FITC)-labeled anti-CD14 (eBiosciences). Appropriate isotype controls (eBiosciences) were also used. Erythrocytes were lysed using FACS lysis solution (BD Biosciences, USA) and the cells were washed with washing buffer followed by centrifugation at 300 × g for 5 min. The cell pellet was then re-suspended in 500 μl of sheath fluid and analysed flow cytometry in FACS caliber. Expression of TLR4, TLR5 and TLR9 on monocytes was quantified in 10 subjects with ins/ins and 10 with ins/del. About 50ul of whole blood was stained with anti-CD14-FITC and anti-TLR4-PE (eBiosciences) or TLR5-PE (Santacruz), expression was analysed as described above by FACS. For TLR9 staining, cells were labeled with anti-TLR9-FITC (eBiosciences) and a membrane marker anti-CD14-PE (eBiosciences), fixed and permeabilized, and analysed by flow cytometry.

### Genotyping of TLR4 (Asp299Gly, Thr399Ile) and TLR1 (I602S) polymorphisms

Genomic DNA of all healthy lab volunteers were analyzed for TLR4 (Asp299Gly, Thr399Ile) and TLR1 (I620S) polymorphisms. Genotyping of TLR4 Asp299Gly polymorphism was carried out by allele specific PCR. Two forward primer (allele specific discrimination at the 3’ end) 5’-CTTAGACTACTACCTCGATGA-3’ (wild), and 5’-CTTAGACTACTACCTCGATGG-3’ (mutant), with a common anti-sense Reverse Primer-5’-TAAGCCTTTTGAGAGATTTGA-3’ were used and PCR was performed as follows: 12 cycles of 10 seconds at 950 C followed by 60 seconds at 650C, followed by 17 cycle of 10 seconds at 950C, 50 seconds at 600C and 30 seconds at 720 C. The final extension was for 7 min at 720C. β-actin primers were taken as internal controls. The amplified products (192bp) were analyzed by electrophoresis on 2.5% agarose gels stained with ethidium bromide. DNA from normal subjects will get amplified with ‘wild type’ primers - heterozygous mutations will be characterized by amplification with both ‘wild type’ as well as ‘mutation’ specific primers. DNA from individuals homozygous for TLR4 mutations will get amplified only with mutant specific primer. TLR4 Thr399Ile polymorphism was detected by PCR amplification of involved region with sense primer 5’-GCTGTTTTCAAAGTGATTTTGGGAGAA-3’ that contains a mismatch at position –3 from the 3’ end of the primer (G instead of C) to create a *HinfI* cleavage site when the mutant allele is amplified. Antisense primer was 5’-CACTCATTTGTTTCAAATTGGAATG-3’. The parameters were an initial denaturation at 95°C for 7 min, followed by 35 cycles: denaturation at 95°C for 30 s, annealing at 62°C for 30 s, and elongation at 72°C for 32 s. The final elongation was at 72°C for 5 min followed for a cooling to 4°C. The amplicon of 147-bp length were digested at 37°C with Hinf I for 4 hrs resulting in fragments that either were cut into two fragments of 122-bp and 25-bp (mutant allele) or were not restricted (wild allele). These fragments were analyzed by electrophoresis on 3% agarose gels stained with ethidium bromide. PCR-RFLP technique was also employed to determine variation in codon 602 of TLR1 gene. A fragment containing exon 4 of TLR1 was amplified from genomic DNA using the following PCR primers: forward, CTTGATCTTCACAGCAATAAAATAAAGAGCATTCC, and reverse, GGCCATGATACACTAGAACACACATCACT. The PCR conditions were an initial denaturation at 95°C for 5 min, followed by 32 cycles: denaturation at 95°C for 30 s, annealing at 60°C for 30 s, and elongation at 72°C for 32 s. The final elongation was at 72°C for 5 min. The amplicon was digested by PstI to discriminate between the TLR1 602 alleles.

### TLR2 GT repeats typing

To amplify a region of about 150 base pairs flanking the microsatellite GT repeat, We performed PCR using the FAM-labeled primer (forword-FAM-5’-TATCCCCATTCATTCGTTCCATT-3’) and a reverse primer (5’-GACCCCCAAGACCCACACC-3’) (Integrated DNA technologies, Iowa USA). The thermal cycler conditions were as follows: preheating at 94°C for 5 minutes, followed by 35 cycles of 94°C for 30 seconds, 58°C for 30 seconds, and 72°C for 30 seconds; and then a final extension of 5 minutes at 72°C. The numbers of GT repeats were determined by sizing the PCR products using an ABI 310 sequencer (Applied Biosystems, Foster City, CA) and Genescan analysis 2.1 software (Applied Biosystems). For all primates, same primer set were used for typing of TLR2 microsatellite GT repeats.

### Isolation and stimulation of monocyte

About 5 ml of whole venous blood were collected from pre-genotyped healthy lab volunteers that include: seven ins/ins individuals, seven subjects of ins/del genotype and five homozygous del/del individuals. All subjects enrolled for functional study with ins/ins, ins/del or del/del genotypes were wild type for TLR4 (Asp299Gly and Thr399Ile) and TLR1 (I602S) polymorphisms. 5 ml of blood was layered onto 5 ml of HISTOPAQUE-1077 (Sigma) and centrifuged at 400 g for 30 min in room temperature. Peripheral blood mononuclear cells were removed from the interface and washed twice (10 min, 250 g) in culture medium before being counted. Cells were resuspended to 10^6^ cells per ml in Dulbecco’s Modified Eagle Medium (DMEM) with 1% antibiotics (Penicillin Streptomycin solution from Sigma) and 10% fetal bovine serum. Cells were placed in a 96 well culture plate (100μl per well) and incubated in a 5% CO_2_ incubator at 37°C for 4 hours. Non adherent cells were removed by rinsing of medium thrice. Adherent cells were stimulated with wide range of TLR2 agonists: Peptidoglycan from Streptomyces species, Methanobacterium species, Saccharomyces cerevisiae, Micrococcus luteus, Bacillus subtilis, and Staphylococcus aureus (100μg/ml), all obtained from Sigma.

Synthetic TLR2 agonist Pam3CSK44 (1ng/ml) from IMGENIX, Vi polysaccharide (20μg/ml) from GlaxoSmithKline and γ- irradiated Mycobacterium tuberculosis (1X10^6^ /ml). LPS was taken as positive control (10μg/ml) (Sigma) and monocytes were stimulated with CpG (10μg/ml), Poly IC (10μg/ml) and flagellin (100μg/ml) as negative control. LPS contamination was minimized by addition of 10μg/ml of polymixin-B to the culture medium except positive control well (LPS 10ug/ml Sigma). After 48 hours of incubation supernatant were collected and stored at −80°C until assays for cytokine.

### Primates DNA

Blood samples of four rhesus monkeys and three baboons (National institute of Immunology, New Delhi) were collected and whole genomic DNA was isolated by DNA isolation kit (Sigma). Fecal samples of five chimpanzees were collected from captivated animals at Nandan Kanan Zoo, Bhubaneswar, Odisha, India. Fecal DNA was extracted by commercial kit according to manufracturer’s instructions (Qiagen). Fecal DNA of 25 Chimpanzees and 8 Gorilla were a kind gift from Prof. B.Hann.

### Multiple sequence alignment

Nucleotide sequence flanking 23bp ins/del polymorphic site of TLR2 gene of all primates whose whole genome sequence data are available in NCBI or Ensembl data base were obtained (*Homo sapiens*, *Pan troglodytes*, *Pongo abelii*, *Macaca mulatta*, *Callithrix jacchus*, *Papio hamadryas*). Recently, Neanderthal genome sequence was reported by sequencing of three Neanderthal bones from the Vindija Cave in Croatia: Vi33.16, Vi33.25, and Vi33.26. The whole genome sequence consists of approximately 50bp short sequence fragments. These sequences were downloaded from database. Our sequencing data (del/del genotype of *Homo sapiens*, eight sequence of *Gorilla gorilla*, four *Macaca mulatta*, three *Papio hamadryas* and five *Pan troglodytes*) were aligned using the ClustalW program. All default parameters of ClustalW program were used.

### P. falciparum malaria patients

*P. falciparum* infected subjects were enrolled at S.C.B. Medical College and Hospital, Cuttack, Odisha. The area is highly endemic for malaria with more than 85% of malaria cases attributed to *P. falciparum*. Patients enrolled in the current study are residents of coastal districts in India which are highly endemic for *P. falciparum* malaria – average Annual parasite index (API) of *P falciparum* was 6.67 (http://nvbdcp.gov.in/Doc/malaria_situation_april12.pdf). Clinically suspected malaria patients were screened for *P. falciparum* infection by Giemsa-stained thick and thin blood smears and immune chromatography test (SD Bio Standard Diagnostics India). Detection of *P. falciparum* was also performed by nested polymerase chain reaction (PCR) (*10*). *P. falciparum* infected individuals were categorized based on WHO guidelines. The inclusion criteria for uncomplicated malaria (UM) cases were symptoms of fever with evidence of *P. falciparum* infection in the blood. Severe malaria (SM) patients were categorized into three groups based on distinct clinical features: 1) Cerebral malaria (CM), 2) Non cerebral severe malaria (NCSM) and 3) Multi-organ-dysfuction (MOD) as described in our earlier reports (*10, 37*). About 5ml of venous blood was collected in EDTA vials from all enrolled patients. Plasma was separated a stored at −80°C until further use.

### C-reactive protein (CRP) and Cytokines (TNF-α, IL1β) assays

Plasma CRP of healthy lab controls and *P. falciparum* infected patients were quantified by quantitative immunoturbidometric assays according to manufacturer’s instructions (Labkit, Merck, Millipore). Enzyme linked immunosorbent assay (ELISA) was employed for quantification of plasma TNF-α from healthy lab controls and falciparum infected patients according to manufracturer instructions (ebiosciences). IL1β levels in culture supernatant were also quantified by bioplex (Biorad) or ELISA kit (ebiosciences).

### Statistical analysis

GraphPad Prism (Version 5.01) was used for all statistical analysis in the study. Clinical characteristics and biochemical analysis of patients were compared by one-way analysis of variance (ANOVA). All cytokines and plasma CRP data were analyzed using student’s t test. Correlation between plasma TNF-α and CRP was analyzed by Spearman’s rank test. Allele and genotype frequencies were calculated by direct counting. Fisher’s exact test was used to analyze distribution of allele and genotype frequencies with in different groups of *P. falciparum* malaria infected patients and association with mortality. In addition, odds ratios (ORs) and 95% confidence intervals (CIs) were calculated. Deviations from the Hardy-Weinberg equilibrium (HWE) were tested using the web-based site http://www.oege.org/software/hwe-mr-calc.shtml. Statistical analysis of P value less than 0.05 was considered as significant.

## Acknowledgments

The authors thank Dr. Krishna Prasad, AIIMS, New Delhi for generous gift of γ-irradiated M. tuberculosis, Dr. Subeer Majumdar, NII, Delhi for samples of normal Rhesus monkeys, Dr. D. K. Panda, Director, Nandankanan zoo, Bhubaneswar for stool samples of Chimpanzees, Prof. Beatrice Hahn, University of Pennsylvania, USA for generous gift of DNA samples of Gorilla and Chimpanzees and Dr. Tushar Vaidya, CCMB, Hyderabad for assisting in analysis of Neanderthal sequences and Prof. Partha Majumder for critically reading the manuscript.

## Author contributions

